# Group A *Streptococcus* strains causing meningitis without distinct invasive phenotype

**DOI:** 10.1101/2023.03.22.533757

**Authors:** Laura Marquardt, Federica Andreoni, Mathilde Boumasmoud, Tiziano A. Schweizer, Dorothea M. Heuberger, Elena Parietti, Sanne Hertegonne, Jana Epprecht, Dario Mattle, Anna K. Raez, Ewerton Marques-Maggio, Reto A. Schuepbach, Barbara Hasse, Srikanth Mairpady-Shambat, Silvio D. Brugger, Annelies S. Zinkernagel

**Author notes:** These authors contributed equally to this manuscript (Laura Marquardt, Federica Andreoni). Corresponding author: Prof. Dr. Dr. med. Annelies Zinkernagel, University Hospital Zurich, Raemistrasse 100, 8091, Zurich.

## Abstract

Group A streptococcal (GAS, aka *Streptococcus pyogenes*) meningitis is a fulminant disease associated with high morbidity and mortality. To elucidate the mechanisms underlying the invasiveness of GAS in meningitis, we compared GAS isolates derived from five cases of meningitis, to otitis and colonizing isolates. We did not observe differences in adherence to and invasion of human brain microvascular endothelial cells, virulence factors activity or barrier disruption. Whole genome sequencing did not reveal particular invasiveness traits. Most patients previously suffered from otitis media suggesting that meningitis likely resulted from a continuous spread of the infection rather than being attributable to changes in pathogen’s virulence.

## Introduction

*Streptococcus pyogenes* or group A *Streptococcus* (GAS) asymptomatically colonizes the human throat and skin. In addition to asymptomatic colonization, GAS causes mild to life-threatening infections, such as necrotizing fasciitis and meningitis. GAS meningitis cases are rare and represent only 1-4% of all invasive GAS cases detected across Europe and the United States [1–3]. GAS meningitis is a fulminant disease associated with high morbidity, including neurologic sequelae, and mortality. It occurs mainly in patients with predisposing factors such as otitis, sinusitis, recent head injury or neurosurgery, presence of a neurosurgical device, cerebrospinal fluid leak, or alcoholism [4, 5]. This suggests extension *per continuitatem* from a neighboring infection such as otitis media, resulting in otogenic meningitis. However, in a third of cases there are no predisposing factors present. The prevalence of the predisposing factors for GAS meningitis in children are acute otitis media and acute mastoiditis (2-3% [6, 7] and 11% [8], respectively). GAS is also responsible for 41% of acute mastoiditis and 14% of acute otitis cases that require hospitalization in children [9, 10] and for 19% of acute otitis and mastoiditis cases requiring hospitalization in adults [11]. The most common host-associated risk factors for invasive GAS infections in adults are crowding, diabetes, cardiac disease, cancer and corticosteroid use [12]. Other risk factors are breaching of the physical barriers skin and mucosa by trauma or preceding viral infections such as varicella [13] and influenza [14, 15].

On the pathogen’s side, GAS components such as the streptococcal virulence factors Streptolysin O (SLO), streptococcal DNases, the IL-8 protease SpyCEP and the M-protein, play key roles in the invasiveness and ability of the pathogen to successfully invade and survive within the host environment [16]. SLO is a pore forming toxin that can lead to inhibition of phagocytosis and cell lysis of target cells [17], the DNase sda1 was shown to degrade neutrophils extracellular traps and to prevent TLR9-dependent recognition, leading to enhanced pathogen spread in the host [18–20], while SpyCEP is involved in degradation of the cytokine IL-8, impairing neutrophils recruitment to the site of infection [21].

The M-protein is a GAS surface protein promoting GAS survival in the host by increasing adherence to host cells [22], escape from opsonophagocytosis [23] and neutrophil activation, leading to inflammation [24]. To date, more than 200 different M-protein types (*emm*-types) have been described, based on the sequence of the variable region of the M-protein. Genotyping methods used for the classification of GAS strains include serotyping of the M-protein as well as Multilocus Sequence Typing (MLST), based on the sequences of seven housekeeping genes [25]. MLST is normally used to study the epidemiology and population genetics of a specific set of isolates, proving particularly useful for the detection and classification of outbreaks. Strains of *emm* type 1 (*emm*-1) and MLST-28 (ST28) or emm-28 ST52 are highly prevalent among GAS meningitis isolates [26].

The aim of this study focused on assessing whether specific pathogen traits distinguish GAS meningitis isolates from colonizing GAS isolates or GAS causing otitis and whether these specific traits could be associated with disease severity. We investigated five patients suffering from GAS meningitis, analyzing both host and pathogen characteristics. On the host side, various clinical parameters were analyzed. On the pathogen side, we phenotypically and genetically compared five GAS isolates derived from the abovementioned meningitis cases with ten GAS isolates derived either from otitis patients or asymptomatic colonization. To our knowledge, this is the first study comparing the virulence traits of meningitis isolates with otitis or colonizing ones.

## Material and methods

### Patients and ethics

Five patients diagnosed between 2013 and 2017 with clinically confirmed community-acquired GAS meningitis in Switzerland were retrospectively included in this study (Tab.1). No specific surveillance program for GAS meningitis exists in Switzerland and the cases were selected using a convenience sampling strategy. Otitis isolates were also selected using a convenience sampling strategy while GAS colonizing isolates were selected to reflect the population present in the meningitis and otitis groups (*emm*-1 and *emm*-28 types). This study was approved by the regional committee for medical research ethics (BASEC-ID 2016-00145 and 2017-02225).

### Bacterial strains and cell lines

All meningitis, otitis and colonizing GAS clinical isolates are listed in Tab.S1 and were cultured in THY [Todd-Hewitt broth (BD) plus 2% yeast extract (Oxoid)] at 37°C. Immortalized human brain microvascular endothelial cells (HBMECs) [27] were cultured in Roswell Park Memorial Institute (RPMI) medium supplemented with 10% FBS and L-glutamine, unless specified otherwise.

### Adherence and invasion assays

Adherence and invasion assays were performed as previously described, with minor modifications [28]. HBMECs were seeded in 24 well plates to 2.2*10 cells/well in RPMI medium supplemented with 10% FBS and L-glutamine and incubated at 37°C in the presence of 5% CO_2_ for 48h. Overnight (ON) cultures of the GAS strains were grown at 37°C in a static incubator in 5ml THY. Bacterial cultures were diluted 1:10 in fresh THY medium and grown to an optical density at 600nm (OD_600_) of 0.4, washed once with PBS and resuspended in infection medium (RPMI+L-glutamine+0.4% bovine serum albumin). Prior to infection, HBMECs were washed once with PBS and bacteria, resuspended in 500 ul of infection medium, were added to reach a multiplicity of infection (MOI) of 1 or 10 for adherence and invasion assays respectively. The plates were spun down at 1200g for 5 minutes to and subsequently incubated 37°C in the presence of 5% CO_2_. To assess adherence, infected HBMECs were incubated for 30 minutes, washed six times with PBS to remove unbound bacteria and subsequently lysed using 500 ul of deionized sterile water (dH2O). To assess invasion, infected HBMECs were incubated for 2 hours and washed three times with PBS to remove excess bacteria. 500 ul of infection medium supplemented with penicillin and gentamycin, at a final concentration of 10 ug/ml and 100 ug/ml respectively, were added to each well to kill extracellular bacteria. The plates were then incubated for 2 hours at 37°C in the presence of 5% CO_2_ and the cells were subsequently lysed with 500 ul of dH2O after three washes with PBS. Ten-fold serial dilution of the cell lysates were plated on THY plates to enumerate bacterial colonies after ON incubation at 37°C. The percentage of adherence and invasion were calculated relatively to the initial inoculum.

### Viability assay

Cell viability was assessed as follows. HBMECs and bacteria were prepared for adherence and invasion assays, as described above. After 30 minutes infection followed by six washes with PBS or after the two hours incubation with antibiotics followed by three washes with PBS, HBMECs were harvested using 100 µl of trypsin (Gibco), washed and resuspended in 500 µl PBS. Three wells/condition were pooled and the experiment was carried out in duplicate.

250 µL of the cell suspension were transferred to the wells of a conical 96well plate, spun down at 470g for 5min, washed with Annexin V binding buffer (ThermoFisher) and spun down at 470g for 5min. Dead cells were stained using 40 µl/well of a cocktail of Annexin V – FITC (#640906; Biolegend, 1:50 dilution) and 7 AAD (#420404; Biolegend 1:25 dilution) diluted in Annexin V binding buffer, for the detection of apoptotic and late apoptotic cells. The cells were incubated for 20min at room temperature, 160 µL of Annexin V binding buffer were added and data were acquired using the AttuneNxT flow cytometer (ThermoFisher).

### Virulence factors activity

Activity of the virulence factors SLO, streptococcal DNases and SpyCEP was assessed in the supernatants of exponentially growing GAS as previously described [29]. ON cultures of the various strains were statically grown at 37°C in THY medium. Bacterial cultures were diluted 1:10 in fresh THY medium and grown to an optical density at 600nm (OD_600_) of OD 0.4. After centrifugation, the supernatants were removed and filter sterilized using 0.22 µm filtration membranes.

SLO activity was assessed as follows. 4 mM DTT and 0.0004% Trypan-blue were added to the supernatants, the samples were incubated for ten minutes at room temperature (RT) and two-fold dilutions were made in PBS in a 96 well plate. PBS and dH_2_O alone were used respectively as negative and positive controls. Erythrocytes were diluted to 2% v/v in PBS and added to the diluted supernatants (1 part erythrocytes, 4 parts supernatant) or PBS or water for no lysis and full lysis controls, respectively. Samples were incubated at 37°C for 30 minutes and spun down at 3000g for five minutes. The OD_541_ of the supernatants, directly proportional to the activity of SLO, was used as a readout.

Streptococcal DNase activity was assessed as follows. 5 ul of bacterial supernatant or 5 ul of THY (negative control) were mixed with 5 ul of 10X DNase buffer (50 mM CaCl2, 500 mM Tris, pH 7.9) and 5ul of calf thymus DNA (1mg/ml, Sigma) in a total volume of 50 ul and incubated at 37°C for five minutes. The reaction was stopped by adding 8 ul of 0.5M EDTA and the samples were loaded on a 0.8% agarose gel to assess DNA degradation. Streptococcal DNase activity was evaluated with a score corresponding to no activity (0=no degradation of calf thymus DNA), intermediate activity (1=partial degradation of calf thymus DNA) and high activity (2=complete degradation of calf thymus DNA).

SpyCEP activity was assessed by measuring the degradation of the cytokine IL-8. Bacterial supernatants were incubated with 1ng/ul IL-8 for 16 hours at 37°C after which an IL-8 ELISA was carried out according to the manufacturer’s instruction (R&D). THY was used as a negative control. SpyCEP activity is depicted as the percentage of IL-8 cleavage compared to the negative control (medium only).

### Electric cell-substrate impedance sensing (ECIS®) assay

ECIS® was performed with the ECIS® Z-Theta instrument (Applied Biophysics) to monitor the effects of the GAS strains on human brain microvascular endothelial cells (HBMECs) [27] barrier integrity. Before seeding, the wells of the arrays (ECIS 8W10E+, IBIDI) were washed twice with sterile, nuclease-free dH_2_O, coated with 5 µg/ml collagen IV for 1h at room temperature (RT), washed three times with dH_2_O and incubated with 10 mM L-cysteine for 2h at RT. The wells were washed two times with dH_2_O before 200 µl of culture medium were added and the assay started (37°C, 5% CO2). After 30 min, 200µl culture medium containing 10 HBMECs were seeded per well. After 22 h incubation 50 µl of medium were removed and 2h later HBMECs were infected with log phase GAS strains resuspended in DMEM+10%FCS at an MOI of 10. DMEM+10%FCS alone was used as a control. After 1.5h penicillin (10 µg/ml) and gentamycin (100 µg/ml) were added. Barrier integrity was monitored at 4000 Hz from seeding to 70h post infection. Resistance data were normalized to the time of infection.

### Bacterial genomics

Whole genome sequencing and genomic data analysis were performed as previously described [30]. Briefly, from 150 bp paired end reads*, de novo* assemblies were generated with SPAdes v.3.10 [30] and annotated with Prokka v.1.13 [31]. A pangenome was constructed with Roary v.3.12 [32] and the resulting core-gene alignment was used to build a maximum likelihood phylogenetic tree with Fasttree v.2.1.10 [33]. *Emm*-typing was performed *in-silico* by querying assemblies against the curated CDC database [34, 35] with blastn v.2.9.0. Virulence genes were identified by querying the assemblies against the full VFDB database, i.e. VFDB_setB_nt.fasta, downloaded on the 04.08.23 [36], using ABRIcate v.0.5 (minimum identity and coverage: 85%). ENA accession number: PRJEB57816.

### Statistics

Three biological replicates were carried out for each strain, except for HBMECs viability and ECIS assessment where two biological replicates were carried out for each strain. The average of the biological replicates for each strain is represented on the graphs as a single data point. The average of these values represents the final value depicted on the graphs. Error bars represent standard deviations. The Mann-Whitney test was used to assess statistical significance for all assays apart from HBMECs viability data that were analyzed using the Kruskal-Wallis test.

## Results

### Cases description

Five GAS meningitis cases, occurring between 2013 and 2017 in Switzerland, were included in this study and the corresponding GAS isolates were retrospectively retrieved for analysis. The median age of the patients was 64 years (42 to 79 years) and 60% (n=3) were female. The mortality rate was 20% (n=1) while 2 out of 6 patients (33%) experienced neurological sequelae (patients 3 and 4), which included slight hearing loss, vertigo and slight paresis of the limbs. The median duration of the hospital stay was 17 days (3 to 29 days) and 4 patients needed intensive care treatment with endotracheal intubation. Relevant clinical and laboratory findings are depicted in table 1. Mastoiditis was the underlying diagnosis in three cases. During the course of the disease one patient had septic thrombosis and one developed a brain abscess (Fig.1A-1B). Surgical treatment such as antrotomy, mastoidectomy, or burr-hole trepanation was required in three patients. In patient 4, histological analysis from the surgically-removed mastoid tissue showed necrotic tissue and individual mucosal fragments covered by simple cuboid epithelium with severe acute and chronic inflammation and showing extracellular accumulation of bacterial cells (Fig.1C-1D).

**Figure 1.**
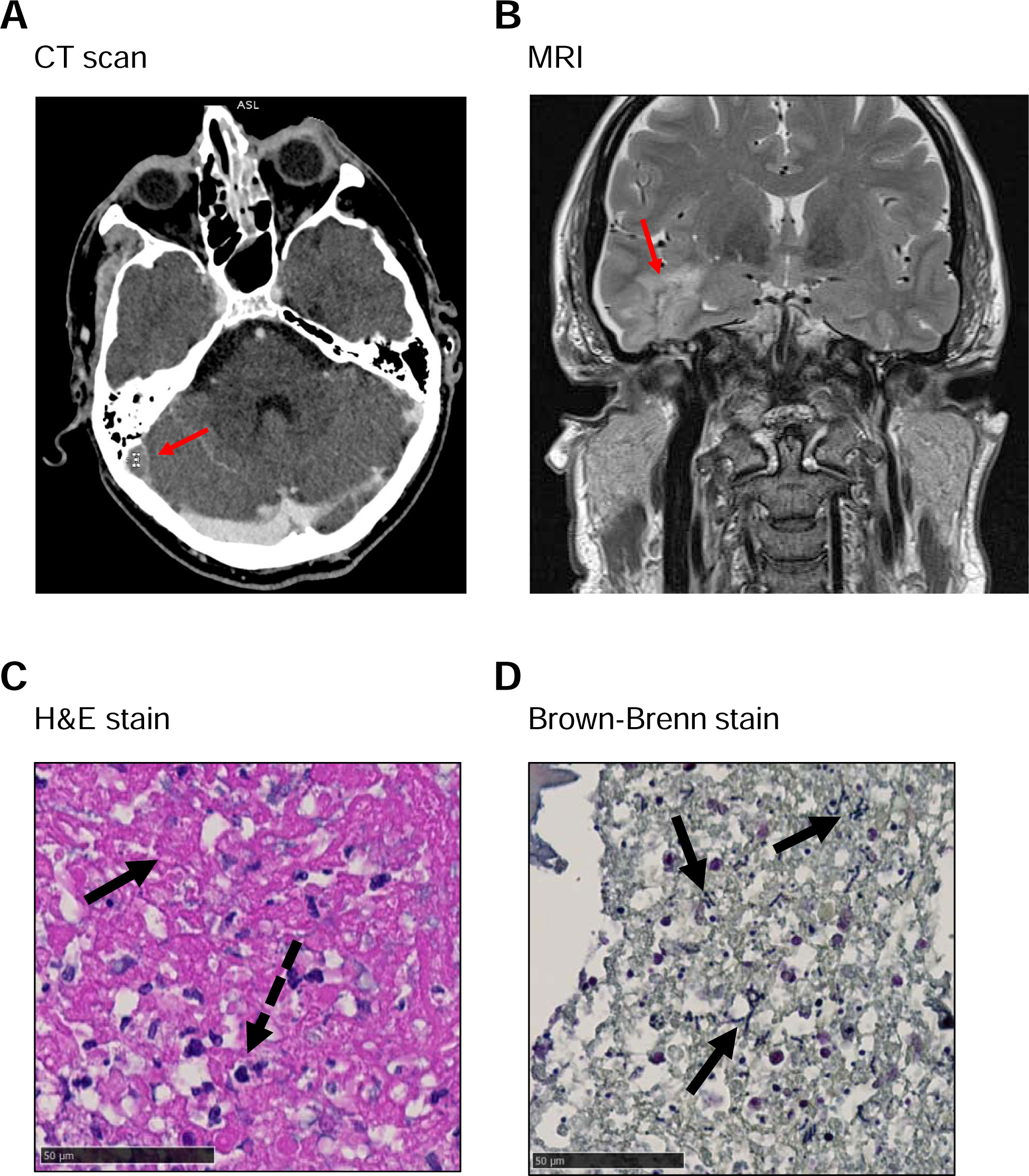
Patients scans and histology results. **A)** CT-Scan: septic thrombosis with mastoiditis and otitis media on the right side. The red arrow indicates thrombosis **B)** MRI-Scan: T2 sequence showing penetration of lower temporal lobe with abscess formation. The red arrow indicates an abscess **C)** HE Staining of mastoid tissue: the black arrow indicates necrotic mastoid tissue, the dashed black arrow indicates mixed neutrocytic and lymphocytic infiltration, foam cells and cholesterol crystals (not depicted) indicating chronic inflammation. **D)** Brown-Brenn staining of mastoid tissue: The arrows indicates accumulation of extracellular bacteria in chains.

**Table 1.** Patients description. Clinical findings, complications and treatment regimen of the patients included in this study. Abbreviations: CSF=cerebrospinal fluid; d=day; ICU=intensive care unit.

| Patient ID<br>Sex<br>Age(y)<br>Isolate Nr. | Associated health conditions | Clinical presentation | Complications | Site of GAS identification | Antibiotic treatment regimen | Death |
| --- | --- | --- | --- | --- | --- | --- |
| 1<br>F<br>73<br>CI1224 | • Perforated otitis media<br>• Diabetes<br>• Hypothyreosis<br>• Depression | • Altered vigilance and mental state<br>• Vertigo<br>• Hypothermia<br>• Hypotension<br>• Acute abdomen | • Partial septic sinus transversus thrombosis<br>• Coma<br>• Irreversible hypoxic brain damage<br>• ICU involvement | CSF<br>Blood<br>Ear swab | 3d ceftriaxon<br>3d amoxicillin<br>3d clindamycin | yes |
| 2<br>M<br>55<br>CI1271 | • Otitis media<br>• Depression | • Altered vigilance<br>• Meningism<br>• Slight one-sided arm paralysis<br>• Otalgia<br>• Diarrhea<br>• Nausea | • Coma<br>• Mastoiditis<br>• Cerebral abscess<br>• ICU involvement | Blood<br>Mastoid tissue | 2d ceftriaxon<br>1d amoxicillin<br>Escalation to:<br>18d penicillin<br>5d metronidazole<br>5d clindamycin<br>+ Surgical intervention | no |
| 3<br>F<br>64<br>CI1293 | • Bicuspid aortic valve | • Acute reduction of general condition<br>• Meningism<br>• Right sided paresis of upper and lower limbs<br>• Right sided tongue deviation<br>• Anisocoria<br>• Petechial bleeds | • Thrombocytopenia<br>• Acute kidney failure<br>• Aortitis<br>• Nosocomial Influenza infection | CSF<br>Blood | Initially ceftriaxon then change to penicillin due to elevated transaminases, total 18d<br>3d gentamycin<br>15d steroids<br>10d vancomycin<br>10d clindamycin | no |
| 4<br>F<br>79<br>CI407 | • Otitis media (perforated)<br>• Coronary heart disease<br>• Pre-diabetic metabolism | • High temperatures<br>• Vertigo<br>• Altered mental state and vigilance | • Tegmen tympani penetration<br>• Mastoiditis<br>• Labyrinthitis<br>• Coma<br>• Long-term one sided deafness<br>• ICU involvement | Mastoid tissue | Initially 1d ceftriaxon, then escalation to meropeneme for 2d, then back to ceftriaxone for 20 days<br>Initially 2d acyclovir<br>1d amoxicillin<br>4d mephamesone<br>+ Surgical intervention | no |
| 5<br>M<br>42<br>CI543 | • Otitis media<br>• Bullous<br>• Myringitis with insertion of tympanostomy tube | • One day after insertion of tympanostomy tube acute reduction of vigilance<br>• Altered mental state and meningism | • Mastoiditis<br>• Subdural empyema<br>• Cerebritis<br>• Ventrikulitis<br>• Coma<br>• ICU involvement | Blood<br>CSF<br>Mastoid tissue | 9d ceftriaxon, then escalation to 18d ceftazidime and de-escalation back to ceftriaxone for another 35d<br>2d amoxicillin + clavulanic acid<br>2d acyclovir<br>2d metronidazol 2d, pause 3d, then another 2d<br>9d intrathecal gentamycin<br>9d intrathecal vancomycin<br>+ Surgical intervention | no |

### Bacterial virulence

The assessment of adherence to and invasion of HBMECs, the activity of the streptococcal virulence factors DNases, SLO and SpyCEP as well as barrier integrity of HBMECs upon GAS challenge was carried out on meningitis, otitis and colonizing clinical isolates. No significant difference between adherence and invasion behavior to HBMECs upon infection with meningitis, otitis or colonizing GAS isolates was found (Fig.2A-2B) and the toxicity of the isolates toward HBMECs during infection did not vary according to their sampling location (Fig.S1A-S1B). Furthermore, no differences in virulence factors activity or the isolates potential to disrupt the barrier formed by HBMECs among the three groups were observed (Fig.2C-2F). A trend, albeit not significant, towards reduced SpyCEP activity was observed in the meningitis group (Fig.2D).

**Figure 2.**
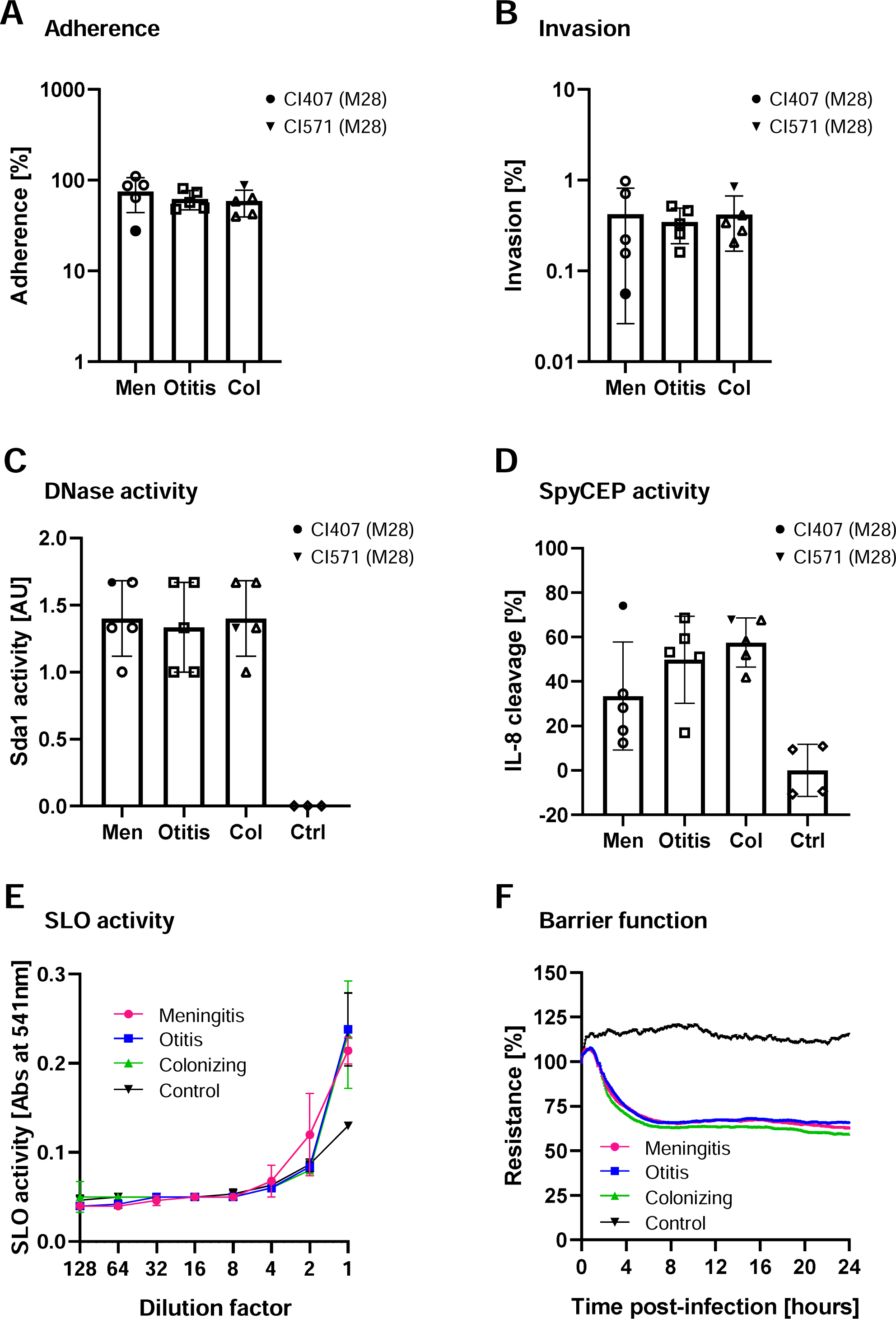
Bacterial strains virulence determinants. **A)** Adherence to and **B)** invasion of human brain vascular endothelial cells (HBMECs) were assessed after 30 minutes (MOI 1) or two hours (MOI 10) of infection, respectively. Adherence and invasion percentages were calculated based on the initial bacterial inoculum used to infect HBMECs. Bacterial adherence above 100% occurred in one case and could be explained by increased initial bacterial growth or by disruption of long bacterial chains or clumps due to sheer pipetting forces during the eukaryotic cells lysis step. Bacterial virulence factors activity of **C)** Streptococcal DNases, **D)** the IL-8 protease SpyCEP and **E)** the pore-forming toxin SLO were assessed in supernatants of exponentially growing GAS. **F)** HBMECs barrier function disruption was carried with the ECIS® Z-Theta instrument (Applied Biophysics), measuring the impedance generated by the cell barrier in the presence or absence of bacteria. Control experiments were carried out using medium only. **A, B, C, D)** The average of the biological replicates for each strain is depicted as a single data point on the graphs, so that each data point represents a single strain. The two strains indicated in the figure legend with a black symbol (CI407 and CI571) are the only strains of *emm*-type 28 (M28), as opposed to all other strains that are *emm-*type 1 (M1). **A, B, C, D, E)** Three biological replicates were carried out for each strain. The average values for strains belonging to a single group of isolates (meningitis, otitis or colonizing) is depicted in the graphs and the error bars represent the standard deviation. **F)** Two biological replicates were carried out for each strain and the average of the strains belonging to the same group is depicted on the graph. Control=medium only. No significant differences in adherence, invasion, virulence factors activity or barrier disruption was observed among the different groups of isolates. A significant difference was found between the respective controls and meningitis, otitis or colonizing isolates for DNase (p=0.0159, 0.0159 and 0.0159, respectively) and SpyCEP (p=0.0179, 0.0357 and 0.0179, respectively) activity. Statistical significance was assessed using the Mann-Whitney test. Men=meningitis, Col=colonizing, Ctrl=control.

### Genotyping

Typing of meningitis and otitis isolates showed a high prevalence of emm-1/ST28 (n=4/5 and n=5/5, respectively, Tab.S1). Since one meningitis clinical isolate belonged to the emm-28/ST52, we chose to include four emm-1/ST28 and one emm-28/ST52 in the colonizing isolates group. Only one meningitis clinical isolate belonged to the *emm*-28/ST52 and one colonizing isolate to the *emm*-28/ST458. A core genome phylogeny revealed that, within the M1 clade, the meningitis isolates did not form a monophyletic group. (Fig.S2A-S2B). The presence of genes encoding virulence factors was analyzed and this showed no specific pattern for meningitis isolates when compared to otitis or colonizing ones (Fig. S2C). We additionally checked for the presence of mutations in the GAS CovRS regulatory genes that may lead to upregulated virulence factors activity (Tab.S2) [37]. No mutations were found in CovR, while we found CovS mutations in two of the meningitis isolates (CI1224, *emm*-1 and CI407, *emm*-28) and in one of the colonizing ones (CI571, *emm*-28). An in-frame deletion of the proline in position 16 was found in CI1224. CI407 presented a deletion of the first 46 amino acids as well as non-synonymous mutations in position 47 (Leu-Met), 226 (Glu-Gly) and 332 (Val-Glu). Isolate CI571 had the same non-synonymous mutations as CI407 in position 226 (Glu-Gly) and 332 (Val-Glu), common to the emm-28 lineage [38]. However, these mutations did not result in a reduced CovS function or increased virulence factors activity.

## Discussion

We found no particular phenotypic or genotypic characteristics in the five GAS isolates causing meningitis as compared to otitis or colonizing isolates. The only common denominator was preceding otitis and mastoiditis on the host’s side. GAS meningitis is an extremely rare condition, representing only the 0.2%-0.9% of cases of bacterial meningitis in the United States and Brazil [39–41] and 1-4% of invasive GAS cases in Europe and the United States [1–3]. A report from Switzerland described nine cases of GAS meningitis registered between 1983 and 1999 across all major Swiss hospitals, again emphasizing the rarity of this disease [42]. A recent report showed an increase in the number of GAS meningitis case in the Netherlands in 2022-2023, compared with the previous 40 years, with emm-1 being the most represented M-protein variant [43].

Meningitis occurs either by hematogenous spread or by direct spread of bacteria from a neighboring infection. Hematogenous spread normally occurs when colonizing bacteria breach the nasopharyngeal mucosa, reach the blood [44] and subsequently cross the blood-brain barrier. It was suggested that spread from the upper respiratory tract’s mucosa may be due to an altered balance in the host microbe population, possibly favoring GAS number increase and subsequent spread [5]. This interesting hypothesis was however never tested and remains a speculation. On the other hand, direct spread to the brain can also occur *per continuitatem* from infected adjacent anatomic structures, such as the ear or the mastoid bone [44].

We searched for common traits in the clinical presentation of our patients and characteristics shared by the meningitis clinical isolates, as opposed to otitis and colonizing ones, to find possible explanations for the spreading of GAS to the brain. In our patients’ cohort, the presence of mastoiditis in four patients and thrombosis in one supports the idea of direct spread through bone erosion from neighboring oto-mastoiditis and osteo-thrombophlebitis. Moreover otitis, a condition often associated with GAS meningitis [5], was present in the majority of cases suggesting continuous spread as a likely cause of infection. In contrast, hematogenous spread was less common and a plausible cause of infection for patient 3 only. There is no way to determine whether the presence of bacteria in the blood in patients 1, 2 and 5 occurred before or after spread to the central nervous system. However, given the infection focus and the evidence pointing to continuous spread from the ear and mastoid bone as one of the main causes of GAS meningitis, we postulate that hematogenous spread, In the case of patients 1, 2 and 5, was not the cause of meningitis.

The presence of a local focus of infection may explain the high surgery rate in our GAS meningitis cohort, which has been reported previously. In a nationwide cohort study in the Netherlands, 46% of GAS meningitis patients required surgical intervention, while this remains an absolute rarity only described in case reports for the more common meningitis pathogens (Pneumococcus, Meningococcus) [4, 45].

On the pathogen side, no common features explaining bacterial spread to the brain were found among the meningitis isolates, as compared to otitis and colonizing ones. This points to host rather than pathogen factors playing a role in facilitating bacterial invasion, as previously described for *S. aureus* [30]. Adherence and invasion assays towards HBMECs confirmed the absence of a particular tropism towards brain microvascular tissue for the meningitis isolates, excluding an advantage in crossing or damaging the blood brain barrier, as compared to otitis and colonizing ones, in the tested settings. Moreover, virulence factors activity did not significantly vary among the three groups of isolates. The *emm*-type of the strains did not generally seem to influence adherence, invasion or virulence factors activity, although the number of strains and *emm*-types tested is too little to be able to make a conclusive statement. A trend towards decreased SpyCEp activity was observed in the meningitis group with the M28 strain CI407 acting as an outlier and presenting a level of activity remarkably higher than the other meningitis isolates. CI407 was characterized by a pronounced deletion of the first portion of the CovS protein, a key regulator of virulence factors activity in GAS, as well as by the presence of three non-synonymous mutations (Tab.S2), as compared to the sequence of the CovS protein of GAS M1T1 5448 [37]. Normally a complete loss of CovS function, as for the GAS M1T1 5448 AP strain, would lead to a complete IL-8 degradation leading to 100% cleavage [29], which we did not observe. SLO and DNase activities were also identical to the non-mutated strains. A partial loss of CovS functionality could be postulated in this case [46], although the mutations in CI407 mainly affect the N-terminal portion of the CovS protein where no functional domains are present [18, 47]. Further investigations involving a larger group of clinical isolates, including different *emm*-types, is needed to address the question whether a decreased IL-8 degradation potential might facilitate invasion of the blood brain barrier. Isolates from the *emm*-1 and *emm*-28 lineages represented both in the meningitis and colonizing groups, behaved very similarly in terms of virulence factors activity.

In accordance with previous studies stating an association of *emm*-1/ST28 with GAS meningitis [26], 4/5 meningitis isolates belonged to the *emm*-1/ST28 lineage. The fifth isolate belonged to the *emm*-28 lineage, which is another prevalent *emm*-type previously described in GAS meningitis [26] and, together with *emm*-1, among the most representative *emm*-types for invasive GAS infections across Europe and nord America [48]. All otitis and colonizing isolates belonged either to the *emm*-1 or the *emm*-28 lineages as well. A prevalence of the emm-1 linage in acute otitis media in children (around 30%) was previously reported [49]. The serotype of colonizing isolates was however specifically selected to reflect the serotypes present in the meningitis and otitis groups. It cannot be excluded that the overrepresentation of *emm*-1 serotypes in meningitis and otitis might reflect the global success and dominance of *emm*-1 strains causing GAS invasive infections [48]. The presence or absence of GAS virulence determinants did not reveal a specific pattern for meningitis isolates when compared to otitis and colonizing ones.

GAS meningitis occurs very rarely, leading to the availability of a small number of cases that could be included in this study. Nevertheless, despite the small sample size, our data suggest that the most common pathophysiological cause in this patients’ cohort was local spread from a site of bacterial infection in neighboring tissues, with hematogenous spread a possible explanation in one case.

In the absence of distinct virulence determinants characterizing the meningitis isolates, we conclude that, in this particular cohort, host factors such as breach of the mucosa or local infections play a preponderant role in the transition from colonization to invasion. These results call for future studies including a higher number of cases and clinical isolates and a more comprehensive phenotypic and genotypic investigation to support our conclusion.

## Author contributions

LM: Experimental design, acquisition, analysis and interpretation of data, writing of the manuscript.

FA: Experimental design, acquisition, analysis and interpretation of data, writing of the manuscript.

MB: Acquisition, analysis and interpretation of data, writing of the manuscript

TAS: Experimental design, data acquisition, data analysis and critical reading of the manuscript

DMH: Experimental design, data acquisition, data analysis

EP: Experimental design, data acquisition, data analysis

SH: Experimental design, data acquisition, data analysis

JE: Data acquisition, critical reading of the manuscript

DM: Data acquisition

AKR: Data acquisition

EMM: Histological analysis

RAS: Funding

BH: Funding

SMS: Experimental design, funding, critical reading of the manuscript

SDB: Conceptualization, funding, critical reading of the manuscript

ASZ: Conceptualization, funding, critical reading of the manuscript

