## Supplemental material for "Group A *Streptococcus* strains causing meningitis without distinct invasive phenotype"

|  |  |  |  |  |  |  |  | **MIC (mg/l)** | | | |
| --- | --- | --- | --- | --- | --- | --- | --- | --- | --- | --- | --- |
| **Group** | **Patient ID** | **Strain ID** | ***emm*-type** | **ST-type** | ***covR*** | ***covS*** | **Origin** | **PEN** | **AMX** | **CLI** | **CMX** |
| **Meningitis** | 1 | CI1224 | M1 | 28 | wt | mut | Blood, ear swab, liquor | 0.008 | 0.004 | 0.004 | 0.002 |
|  | 2 | CI1296 | M1 | 28 | wt | wt | Blood | 0.008 | 0.004 | 0.002 | 0.002 |
|  | 3 | CI407 | M28 | 52 | wt | wt | Blood | 0.008 | 0.004 | 0.002 | 0.002 |
|  | 4 | CI543 | M1 | 28 | wt | wt | Tissue | 0.008 | 0.004 | 0.002 | 0.015 |
|  | 5 | CI1271 | M1 | 28 | wt | wt | Blood, mastoid tissue | 0.008 | 0.004 | 0.06 | 0.015 |
| **Otitis** | - | CI401 | M1 | 28 | wt | mut | Inner/middle ear | - | - | - | - |
|  | - | CI505 | M1 | 28 | wt | wt | Inner/middle ear | - | - | - | - |
|  | - | CI569 | M1 | 28 | wt | wt | Inner/middle ear | - | - | - | - |
|  | - | CI741 | M1 | 28 | wt | wt | Middle ear | - | - | - | - |
|  | - | CI347 | M1 | 28 | wt | wt | Inner/middle ear | - | - | - | - |
| **Colonizing** | - | CI571 | M28 | 458 | wt | mut | Throat swab | - | - | - | - |
|  | - | CI5011 | M1 | 28 | wt | wt | Vaginal swab | - | - | - | - |
|  | - | CI381 | M1 | 28 | wt | wt | Throat swab | - | - | - | - |
|  | - | CI387 | M1 | 28 | wt | wt | Vaginal swab | - | - | - | - |
|  | - | CI400 | M1 | 28 | wt | wt | Vaginal swab | - | - | - | - |

**Table S1 – Clinical isolates description**. Clinical isolates’ *emm*-type, ST-type, origin, *covRS* genotype and MIC for the antibiotics penicillin (PEN), amoxicillin (AMX), clindamycin (CLI) and co-trimoxazol (CMX) are shown in the table. Details for the mutations found in CovR are depicted in table S2.

| **Group** | **Strain ID** | ***emm*-type** | **Mutations in CovS** |
| --- | --- | --- | --- |
| **MEN** | **CI1224** | M1 | In-frame deletion of Proline in position 16 |
|  | **CI407** | M28 | Deletion of the first 46 amino acids |
|  |  |  | Non-synonymous mutation in position 47 (Leucine to Methionine) |
|  |  |  | Non-synonymous mutation in position 226 (Glutamic acid to Glycine)* |
|  |  |  | Non-synonymous mutation in position 332 (Valine to Glutamic acid)* |
| **COL** | **CI571** | M28 | Non-synonymous mutation in position 226 (Glutamic acid to Glycine)* |
|  |  |  | Non-synonymous mutation in position 332 (Valine to Glutamic acid)* |

**Table S2 – CovS mutations**. The mutations found in the CovS protein are listed above. Mutations were found in CI1224, CI407 and CI571 belonging respectively to the meningitis (CI1224 and CI407) and colonizing (CI571) groups. * These non-synonymous mutations are common to the *emm*-28 type strains. The position specified in the table refers to the CovS protein sequence of the GAS M1T1 5448 GAS strain [1].

**Supplementary figure legends**

**Figure S1 – HBMECs viability during adherence and invasion assays.** The viability of HBMECs was assessed by flow cytometry after staining with Annexin V and 7 AAD, upon bacterial exposure for **A)** 30 minutes (adherence, MOI 1) or **B)** two hours followed by two hours incubation with antibiotics (invasion, MOI 10), according to the adherence and invasion assays protocol. The two strains indicated in the figure legend with a black symbol (CI407 and CI571) are the only strains of emm-type 28 (M28), as opposed to all other strains that are emm-type 1 (M1). Two biological replicates were carried out per strain, per condition. Statistical significance was assessed using the Kruskal-Wallis test.

**Figure S2 – Phylogenetic tree and virulence determinants**. **A and B)** Maximum-likelihood phylogenetic tree based on the core genes alignment, where the color and shape of the tips reflect source of isolation. **A)** Full tree, where the clade of the M1 isolates is collapsed (grey square). CI407 and CI571 are of emm-type 28 and cluster further away from all other isolates of emm-type 1. **B)** Zoom in the clade of the M1 isolates. The scale-bars reflect genetic distance in substitutions/site. Since the total number of variant sites was 12’157, 1 corresponds to 12’157 substitutions and 0.001 corresponds to about 12 substitutions. **C)** Virulence determinants differentially expressed among the clinical isolates genomes (presence: black, absence: white). **EsaG**=Type VII secretion system; **SpeK**=streptococcal pyrogenic exotoxin; **SlaA**=prophage-encoded extracellular phospholipase A2; **SpeC**=Streptococcal pyrogenic exotoxin; **mf2**=nuclease; **enn**=M-family proteins; **mrp**=M-family proteins; **SbfI/PrtF1**=fibronectin binding protein; **SfbII/sof**=serum opacity factor; **sfbX**=fibronectin binding protein; **prtF2**= fibronectin binding protein; **RfbA**= putative enzymes for biosynthesis of the polysaccharide side chains; **M28_RS06795**=gene complement ; **mf3**=nuclease; **SclB**= putative hypervariable surface protein; **Sda**=nuclease; **SpeA**= streptococcal pyrogenic exotoxin; **LepA**=signal peptidase; **FctA**=pilus structure; **SrtC1**=pilum-specific sortase ; **FctB**=pilus structure.

Virulence determinants present in all clinical isolates (only cited in the figure legend): **Fbp54**=Fibronectin binding protein; **Grab**=α2macroglobulin-binding protein; **IdeS/Mac**=IgG-degrading enzyme; **HasA**=Hyaluroninc acid capsule; **HasB**=Hyaluroninc acid capsule; **HasC**=Hyaluroninc acid capsule; **emm**=M protein; **mf/spd**=nuclease; **SpeB**=Streptococcal pyrogenic exotoxin; **SpeG**=Streptococcal pyrogenic exotoxin; **SpeJ**=Streptococcal pyrogenic exotoxin; **ScpA/ScpB**=Streptococcal C5-alpha peptidase; **Ska**=streptokinase; **Slo**=streptolysin O; **SagA**=Bacteriocin-like toxin; **SmeZ**=Streptococcal mitogenic exotoxin Z; **FbaA**=Fibronectin binding protein; **Imb**=Laminin adhesion protein; **HylA**=Hyaluronidase; **sclA**=collagen-like surface protein; **plr/gapA**=Glyceraldehyde-3-phosphate dehydrogenase; **cpA**=collagen type I-binding protein; **htrA/degP**=serine protease; **tig/ropA**=protease; **endoS**=oligosaccharide-cleaving enzyme; **hylP**=hyaluronidase; **spyA**=membrane-bound ADP-rybosyltransferase, **cfa/cfb**=cyclic AMP factor gene, **rfbB**=dTDP-D-glucose dehydratase; **psaA**=surface adhesin A; **shr**=surface protein; **MGAS10750_RS02400**=UDP-glucose/GDP-mannose dehydrogenase family protein; **galU**=UDP–glucose pyrophosphorylase; **eno**=enolase; **SPY_RS07465**=ABC transporter ATP-binding protein; **SPY_RS07470**=iron ABC transporter permease; **isdE**=ABC transporter substrate-binding protein**; CppA**=CppA family protein; **SPY_RS07480**=heme-binding protein
