## Supplementary figures and images for "Group A *Streptococcus* strains causing meningitis without distinct invasive phenotype"

### Supplemental Figure 1

#### Slide 1
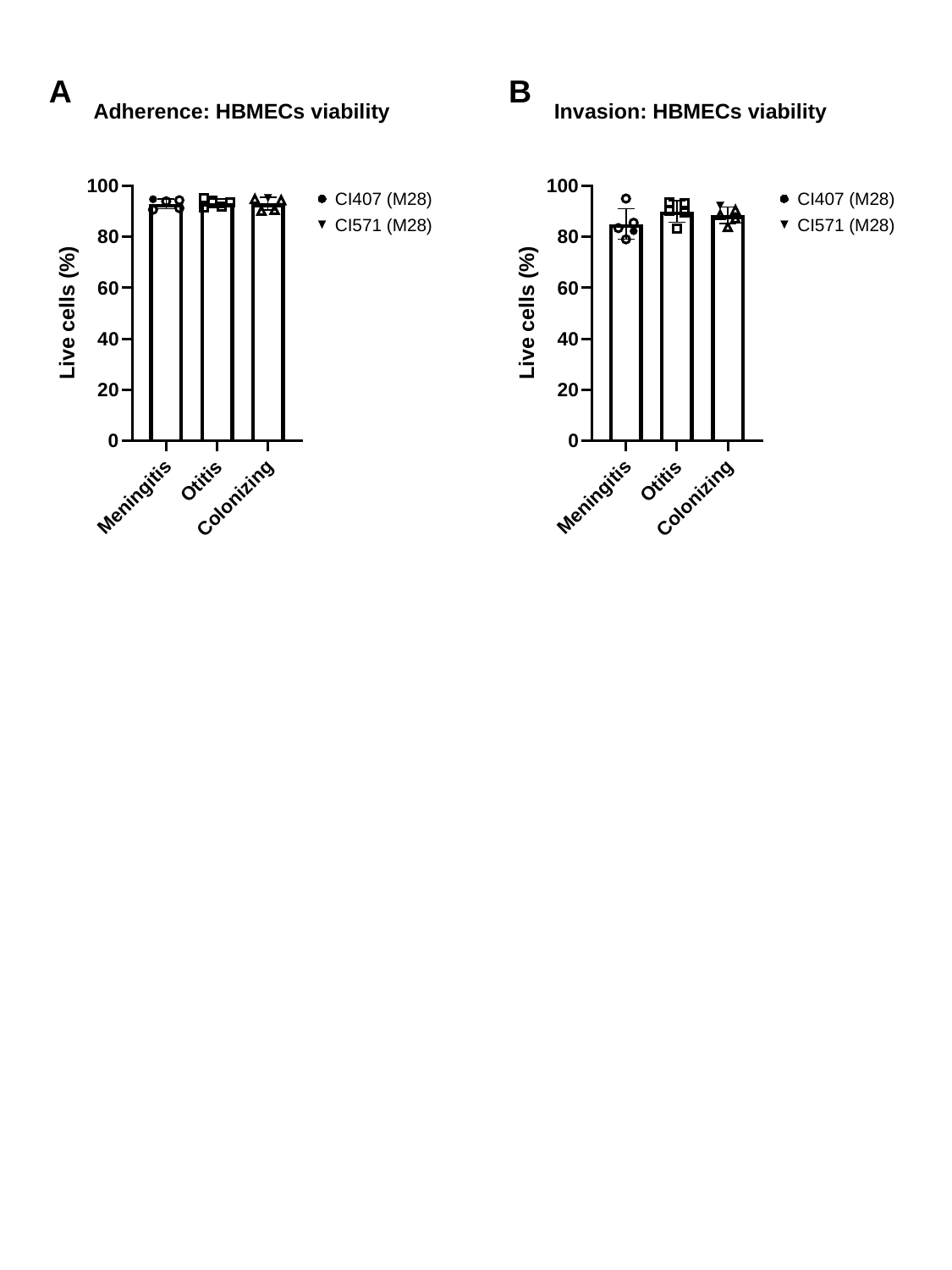

A
B
Adherence: HBMECs viability
Invasion: HBMECs viability
CI407 (M28)
CI571 (M28)
CI407 (M28)
CI571 (M28)

### Supplemental Figure 1

#### Slide 1
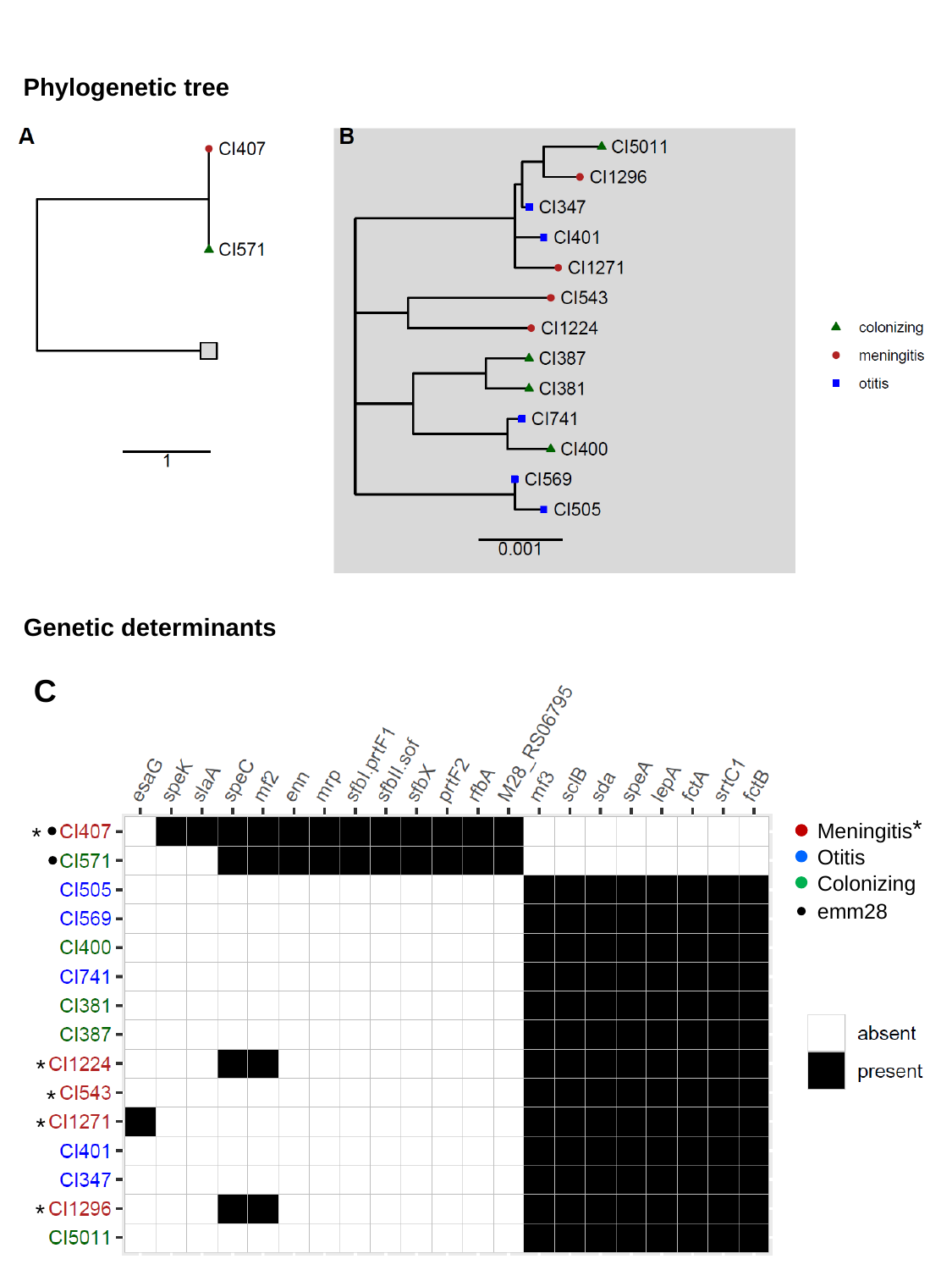

Phylogenetic tree
Genetic determinants
C
*
Meningitis
*
Otitis
Colonizing
emm28
*
*
*
*
